# Seasonal variations in soil fungal communities and co-occurrence networks along an altitudinal gradient in the cold temperate zone of China: A case study on Oakley Mountain

**DOI:** 10.1101/2020.11.17.386136

**Authors:** Li Ji, Yan Zhang, Yuchun Yang, Lixue Yang

## Abstract

The biogeography of soil fungi has attracted much attention in recent years; however, studies on this topic have mainly focused on mid- and low-altitude regions. The seasonal patterns of soil fungal community structure and diversity along altitudinal gradients under the unique climatic conditions at high latitudes remain unclear, which limits our insight into soil microbial interactions and the mechanisms of community assembly. In this study, Illumina MiSeq sequencing was used to investigate the spatiotemporal changes in soil fungal communities along an altitudinal gradient (from 750 m to 1420 m) on Oakley Mountain in the northern Greater Khingan Mountains. Altitude had significant impacts on the relative abundances of the dominant phyla and classes of soil fungi, and the interaction of altitude and season significantly affected the relative abundances of Ascomycota and Basidiomycota. The number of soil fungal taxa and Faith’s phylogenetic diversity (PD) index tended to monotonically decline with increasing elevation. Soil moisture (SM), soil temperature (ST) and pH were the main factors affecting fungal community structure in May, July and September, respectively. The soil dissolved organic carbon (DOC) content significantly shaped the soil fungal community composition along the altitudinal gradient throughout the growing season. Compared to that in May and July, the soil fungal network in September had more nodes and links, a higher average degree and a higher average clustering coefficient. The nine module nodes in the co-occurrence network were all Ascomycota taxa, and the identities of the keystone taxa of soil fungi in the network showed obvious seasonality. Our results demonstrated that altitude has stronger effects than season on soil fungal community structure and diversity at high latitudes. In addition, the co-occurrence network of soil fungi exhibited obvious seasonal succession, which indicated that the keystone taxa of soil fungi exhibit niche differentiation among seasons.

## 1. Introduction

The characteristics of biotic and abiotic factors along altitudinal gradients change extensively with geographic distance such that these gradients are considered natural experimental platforms for evaluating the ecology and evolution of microbial communities in response to environmental change (Siles and Margesin, 2017). Fungi are eukaryotic microorganisms that play important roles in terrestrial ecosystems and can establish symbiotic relationships with plants. They can act as decomposers of animal and plant residues and thereby promote the carbon cycle in forest soils, provide mineral nutrients for plants, and alleviate the carbon limitation of other soil organisms; thus, they play crucial ecological roles (Tedersoo et al., 2014). As in similar studies of plants, animals and soil bacteria (Bryant et al., 2008; Li et al., 2018; Mccain, 2010; Shen et al., 2019; Shen et al., 2013), some recent studies of soil fungi have focused on altitudinal distribution patterns; however, these studies are often performed in low- and mid-latitude regions (Miyamoto et al., 2014; Qi et al., 2017; Sheng et al., 2019; Zhao et al., 2020). Few such studies have been conducted in cold temperate regions at high latitudes, which limits our insight into soil microbial interactions and the mechanisms of community assembly.

In recent years, the diversity patterns and community assembly processes of unculturable microorganisms in extreme environments have attracted more attention (Cox et al., 2016; Shi et al., 2015). According to a study of global fungal biogeography by Tedersoo et al. (2014), the fungal communities of boreal forests and arctic tundra are highly similar. The extremely cold environment of the cold temperate zone is characterized by high soil organic matter contents and severe climatic conditions (low temperatures, strong winds, snow pack, short frost-free periods, etc.), but it nonetheless contains a large number of microorganisms (Xu et al., 2017), perhaps due to the rapid evolution of these organisms and their high genetic diversity (Li et al., 2015). Jarvis et al. (2015) studied ectomycorrhizal fungi in the Cairngorm Mountains, located at high latitudes in Scotland, and found that soil moisture and temperature significantly affected the community composition but not species richness of fungi along an altitudinal gradient (a small altitude range of 300 m). In contrast, studies in some low- and mid-latitude regions have yielded inconsistent results, with the diversity of fungal communities decreasing monotonically (Bahram et al., 2012; Liu et al., 2015; Lugo et al., 2008; Ni et al., 2018) or showing unimodal (Miyamoto et al., 2014; Wang et al., 2015), humpback (Wang et al., 2015) or no significant patterns with increasing elevation (Shen et al., 2014). Many studies have reported that various factors affect the elevational distribution of fungal communities at the local or regional scale. Yang et al. (2019) investigated the rhizosphere soil of dominant woody plants in mountainous areas in eastern China and found that environmental factors and spatial distance explained 24.1% and 7.2%, respectively, of the variation in the diversity of the soil fungal community. Bahram et al. (2012) found that host plants and climate factors play important roles in the altitudinal distribution of ectomycorrhizal fungal communities in the forests of northern Iran. Within a small-scale altitudinal range of alpine tundra on Changbai Mountain, the soil carbon-to-nitrogen ratio was shown to be highly correlated with fungal community composition and significantly negatively correlated with fungal community diversity (Ni et al., 2018). However, most of the samplings in previous studies were carried out at a single time or within a single season, which limits our insight into the spatiotemporal variations in the soil microbial community. Although some studies have observed variations in microbial community composition over time (Delgadobaquerizo et al., 2019; Shade et al., 2013), little is known about the seasonal changes in microbial communities at the local scale in the cold temperate zone.

Recently, Zhang et al. (2020) pointed out that although the α and β diversity of soil bacterial and fungal communities in wheat fields exhibit obvious seasonal variation and that rapidly changing environmental variables are the main factors driving the seasonal succession of soil microbes. However, the effects of large-scale spatial variables on soil microorganisms are far more important than the effects of time. The succession of soil microbial communities over time has an obvious scale effect (Shade et al., 2013). At smaller time scales, the driving factors of microbial community dynamics are intermittent pulses that trigger rapid microbial responses (Placella et al., 2012). Over a longer period of time, the responses of soil microbial communities to environmental variations show obvious seasonal differences (Regan et al., 2014). Averill et al. (2019) investigated the spatiotemporal changes in fungal communities at continental and global scales and found that the similarity of the soil fungal community showed large interannual differences and that the seasonal differences in fungal communities were equivalent to thousands of observed community succession patterns at the scale of a few kilometers. Environmental variables can explain some of these temporal and spatial effects. In high-latitude regions, seasonal dynamics regulate forest ecosystem functions through plant photosynthesis (such as nutrient absorption and input) and soil freeze-thaw cycles (Scott-Denton et al., 2003). Compared with that in other regions in the world, the carbon cycle in high-latitude forests is more sensitive to global warming (Lal, 2005). Therefore, exploring the seasonal succession of soil fungal communities at different altitudes in cold temperate forests will help us understand and predict ecosystem processes under global climate change.

In natural ecosystems, most microorganisms do not exist as separate individuals but form a complex co-occurrence network through direct or indirect interactions (Hallam and Mccutcheon, 2015). The interactions among microorganisms have a greater impact than environmental factors on community structure (Gokul et al., 2016), and network analysis can help us fully understand these complex microbial community structures and the interactions among microorganisms (Barberán et al., 2012; Zhou et al., 2011). In the past few years, the topological characteristics of microbial networks based on biogeography have received considerable attention (Ma et al., 2016; Tu et al., 2020; Wan et al., 2020). Microbial network analysis can reveal the species interactions in niche-related communities and how they are affected by environmental factors (Fuhrman, 2009); furthermore, it can reveal the niche overlap of species and the keystone taxonomic groups that play vital roles. However, we still know very little about the microbial networks of seasonal succession (Zhao et al., 2016), which greatly limits our understanding of the spatiotemporal changes that occur in soil microbial communities.

The Greater Khingan Mountains region is the largest forested area and the highest-latitude area in China; this area has undergone one of the most significant temperature increases worldwide in the past century (Tang and Ren, 2005). Larch coniferous forests are an important part of the northern forest ecosystem in this area. The harsh environment in the cold temperate zone is characterized by an average annual temperature lower than −5 °C and a freezing period of more than 200 days per year, which inhibit fungal activity and growth (Hu et al., 2019). As a result, the distribution and interactions of soil fungi are different from those at low and middle latitudes. In the present study, we used the Illumina MiSeq sequencing platform to explore the seasonal changes in soil fungal communities along an altitudinal gradient in the northern Greater Khingan Mountains. We hypothesized the following: (1) Despite being located at high latitudes, the fungal communities along the altitudinal gradient exhibit obvious biogeographical distribution patterns (unimodal or linear), and soil temperature and moisture are the main factors affecting soil fungal variability along the altitudinal gradient; (2) along the altitudinal gradient, seasonal variation has greater impacts than spatial variation on fungal community structure and diversity; and (3) due to the complex, mutually beneficial-competitive relationship between plants and microorganisms and their different nutrient requirements during the growth season, the soil fungal network exhibits obvious seasonal succession, with network structure being more complex and interactions being stronger at the end of the growing season than at the beginning.

## 2. Materials and methods

### 2.1 Study area and sampling

The study area was on Oakley Mountain (51°50′N, 122°01Ε) at the Pioneer Forest Farm of the A’longshan Forestry Bureau in the northern Greater Khingan Mountains in China. This area is an ideal location for investigating the biogeographical patterns of soil microbes along an altitudinal gradient due to the steep topography. This area is characterized by a cold temperate climate with long, cold winters and short, warm summers. The annual mean air temperature is −5.1°C, and the annual mean precipitation is 437.4 mm. Oakley Mountain is the highest mountain in the northern Greater Khingan Mountains and is covered by snow from October to May, i.e., for approximately 7 months each year (Figs. S1 and S2). The soils are mostly Umbri-Gelic Cambosols according to the Chinese taxonomic system, with an average depth of 20 cm (Gong et al., 1999).

Based on the variation in the composition of the vegetation community along the altitudinal gradient, soil samples were collected from the southern slope of Oakley Mountain at six sites representing different elevations (750, 830, 950, 1100, 1300, and 1420 m, called E1~E6). At each site, we created three 20 × 30 m plots, and eight soil cores of 0~10 cm depth were randomly collected and thoroughly mixed to make a composite sample for each plot. The soil samples were collected in May (early growing season), July (mid-growing season) and September (end of the growing season) 2019 (N=54) and immediately transported on ice to the laboratory. We used a button temperature sensor (HOBO H8 Pro, Onset Complete Corp., Bourne, MA, USA) to record the soil temperature (ST) of every sample. The fresh soil samples were sieved through a 2 mm sieve, and visible roots and other residues were removed. Each of the 54 samples was divided into two subsamples: one that was stored at −80 °C for DNA extraction, and one that was stored at 4 °C for the measurement of soil physicochemical properties. Basic information about the sites at the different elevations is provided in Supplementary Table S1.

### 2.2 Soil physicochemical properties

The soil moisture (SM) and bulk density (BD) were measured by the cutting ring method. The soil pH was measured using a pH meter (MT-5000, Shanghai) after shaking a soil water (1:5 w/v) suspension for 30 min. The soil organic carbon (SOC) and total nitrogen (TN) contents in each sample were measured after tableting using a J200 Tandem laser spectroscopic element analyzer (Applied Spectra, Inc., Fremont, CA, USA), and the total phosphorus (P) content was determined colorimetrically with a UV spectrophotometer (TU-1901, Puxi Ltd., Beijing, China) after wet digestion with HClO_4_-H_2_SO_4_. The soil dissolved organic carbon (DOC) content was analyzed using a total organic carbon (TOC) analyzer (Analytik Jena, Multi N/C 3000, Germany), and the soil nitrate (NO_3_^-^-N), ammonium (NH_4_^+^-N), and total dissolved nitrogen (DTN) contents were determined using a continuous flow analytical system (AA3, Seal Co., Germany). The soil dissolved organic nitrogen (DON) was calculated from the soil contents of NO_3_^-^-N, NH_4_^+^-N, and DTN.

### 2.3 DNA extraction and PCR amplification

Soil fungal DNA was extracted from the soil samples using an E.Z.N.A.^®^ Soil DNA Kit (Omega Biotek, Norcross, GA, U.S.) based on the manufacturer’s protocols. The concentration and purity of the DNA were measured with a NanoDrop 2000 UV-vis spectrophotometer (Thermo Scientific, Wilmington, USA). The quality and quantity of the extracted DNA were evaluated with a 1.0% (w/v) agarose gel. The fungal ITS genes were amplified by nested PCR. The primers ITS3F (5′-GCATCGATGAAGAACGCAGC-3′) and ITS4R (5′-TCCTCCGCTTATTGATATGC-3′) were used to amplify the ITS2 region, which was performed with a thermocycler PCR system (GeneAmp 9700, ABI, USA). PCRs were performed in triplicate in a 20 μL mixture composed of 4 μL of 5× FastPfu Buffer, 2 μL of 2.5 mM dNTPs, 0.8 μL of each primer (5 μM), 0.4 μL of FastPfu Polymerase and 10 ng of template DNA. The PCRs were conducted using the following program: 3 min of denaturation at 95 °C; 30 cycles of 30 s at 95 °C, 30 s for annealing at 55 °C, and 45 s for elongation at 72 °C; and a final extension at 72 °C for 10 min (Caporaso et al., 2012).

### 2.4 Illumina MiSeq sequencing and processing of the sequencing data

A 2% agarose gel was used to extract the PCR products, which were then purified with the AxyPrep DNA Gel Extraction Kit (Axygen Biosciences, Union City, CA, USA). Based on the manufacturer’s protocols, the products were quantified using a QuantiFluor™-ST fluorometer (Promega, USA). Subsequently, the amplicons were merged on the Illumina MiSeq platform (Illumina, San Diego, USA) in equimolar amounts and paired-end sequenced (2 × 300 bp) following the standard protocols of Majorbio Bio-Pharm Technology Co., Ltd. (Shanghai, China). We deposited the raw reads into the NCBI Sequence Read Archive (SRA) database (accession number: SRP286329).

Trimmomatic was used to demultiplex the raw fastq files and conduct quality filtering, and the reads were merged with FLASH with the following procedures implemented (Caporaso et al., 2012): (i) The read segments with an average quality score <20 in a 50 bp sliding window were truncated. (ii) Primers were precisely matched, allowing mismatches between two nucleotides, and read segments with ambiguous bases were deleted. (iii) Sequences with an overlap length of more than 10 bp were combined according to their overlapping portion. UPARSE (version 7.1, http://drive5.com/uparse/) was used to cluster the sequences into operational taxonomic units (OTUs) based on 97% similarity (Edgar, 2013), and UCHIME was used to identify chimeric sequences. The taxonomic identities of the gene sequences for each ITS were assigned by BLAST against the UNITE fungal ITS database at a 90% confidence threshold.

### 2.5 Data analysis

The α diversity indices (observed number of OTUs (Sobs), Chao1, Faith’s phylogenetic diversity (PD), and Simpson) of the Illumina MiSeq sequencing data were analyzed with QIIME (Caporaso et al., 2012). The Shapiro-Wilk test and Levene test were used to evaluate the normality of the data and the homogeneity of variance. Principal coordinate analysis (PCoA) of β diversity based on Bray-Curtis distances was conducted with the ‘vegan’ package in R (version 3.6.1) to analyze fungal community similarity. Analysis of similarities (ANOSIM) and permutation multivariate analysis of variance (PERMANOVA) of the Bray-Curtis distances were conducted to test for differences in the properties of the soil fungal communities among different altitudes and seasons. To identify the effects of soil properties on the soil fungal communities, redundancy analysis (RDA) was used to predict the variation in the communities, and a Mantel test with a Monte Carlo simulation consisting of 999 randomizations was performed. The rda function of the ‘vegan’ package and the mantel.rtest function of the ‘ade4’ package in R were used to perform the RDA and Mantel test, respectively (version 3.6.1) (Team, 2013).

Based on random matrix theory (RMT), the molecular ecological network analysis method (http://ieg4.rccc.ou.edu/mena/) was used to construct networks for the different seasons. In most cases, only nodes detected in half or more of the total sample were retained for subsequent network construction. For more information about related theories, algorithms and procedures, please refer to Deng et al. (2012) and Zhou et al. (2011). Spearman rank correlation was used to establish a co-occurrence network among the soil fungal communities. When constructing the network, the same similarity threshold *(St)* was used to ensure that the co-occurrence networks in different seasons could be compared with each other. Then, the same network size and average number of links were used to generate 100 corresponding random networks. The *Z*-test was used to test for differences between the empirical network and the random networks. The intramodule connectivity value *(Zi)* and intermodule connectivity value *(Pi)* for each node were used to identify the keystone species in the network (Deng et al., 2016; Olesen et al., 2007). In this study, we used the following simplified classification and evaluation criteria: (i) peripheral nodes (*Zi*<2.5, *Pi*<0.62), which have only a few links that almost always connect to nodes in their modules; (ii) highly linked connector nodes (*Zi*’≤2.5, *Pi*> 0.62), which have many modules; (iii) module nodes (*Zi*> 2.5, *Pi*≤0.62), which are highly connected to many nodes in their respective modules; and (iv) network nodes (*Zi*>2.5, *Pi*>0.62), which act as both module nodes and connection nodes. To show the results more clearly, Gephi 0.9.2 was used to visualize the co-occurrence networks of the soil fungi (Bastian et al., 2009).

## 3 Results

### 3.1 Soil physicochemical properties along altitudinal gradients

Altitude had significant effects on BD, SM, ST, pH, NH_4_^+^-N, DOC, and DON. BD was the highest at E2 (Table 1). In all seasons, the SM at E5 was significantly higher than that at E2, E3 and E4 (*P*<0.05). With increasing elevation, soil pH and ST showed decreasing trends. In general, the contents of DOC and DON were lower at lower altitudes. There was no significant difference in SOC or TN among the different altitudes (*P*>0.05), whereas soil total P was significantly affected by altitude (*P*<0.05). Season had significant effects on BD, ST, SOC, TN and NO_3_^-^-N. The ST in July was significantly higher than that in May and September (*P*<0.05). The interaction between season and altitude had no significant effect on any of the soil properties.

**Table 1.**
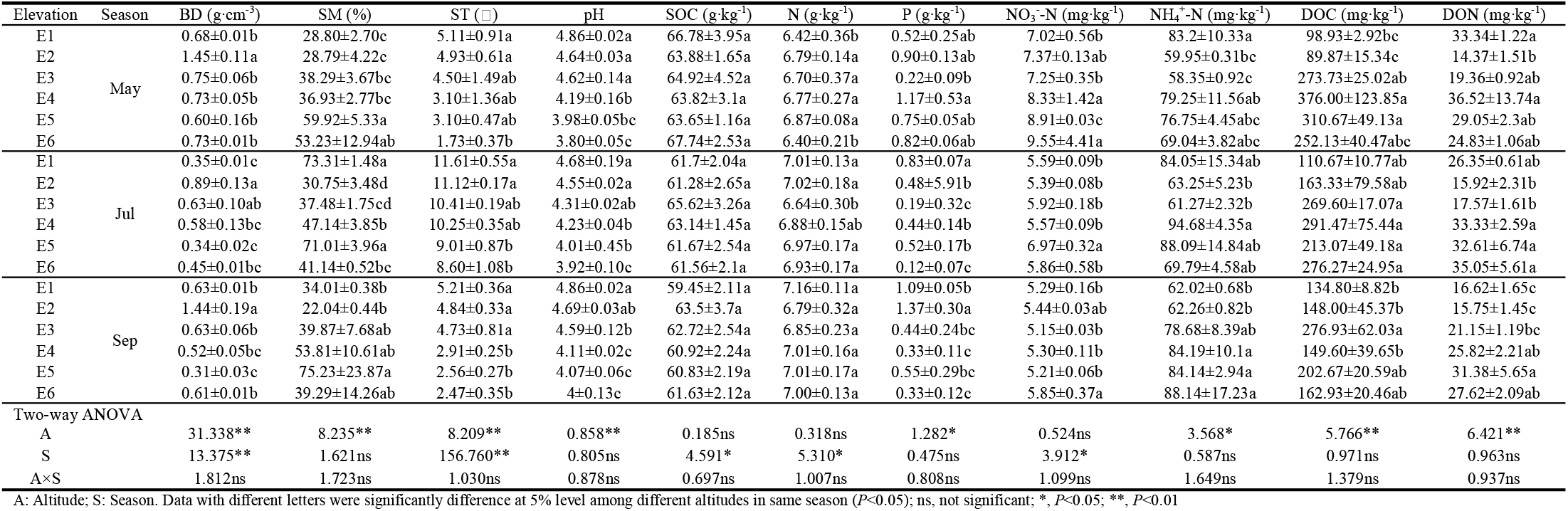
Seasonal dynamics of soil physicochemical property in different altitudes

### 3.2 Soil fungal sequencing characteristics and community composition

Across all soil samples analyzed, 3603426 high-quality soil fungal sequences were obtained by Illumina MiSeq sequencing, representing 1190531, 1177391, and 1235504 sequences from May, July and September, respectively. A total of 42230~74956 soil fungal sequences were obtained per sample (mean = 66730). The average read length, 313 bp, was larger than 99% of Good’s coverage for the ITS gene regions. The rarefaction curves of the genes tended to approach the saturation plateau at 97% sequence similarity for all samples (Fig. S3), which indicated that the sequencing depth was adequate for evaluating the structure and diversity of soil fungi across all samples.

Based on the 97% sequence similarity level, a total of 747 genera and 4106 OTUs were identified, distributed in 16 phyla and 64 classes. Ascomycota (65.8%) and Basidiomycota (31.2%) were the dominant phyla, accounting for 97.0% of the total number of fungal sequences obtained (Fig. 1A). At the highest altitude (E6), the relative abundances of Ascomycota in May and September were lower than the relative abundance in July, at 33.4% and 58.8%, respectively. At the class level, Agaricomycetes (29.2%), Leotiomycetes (25.4%) and Eurotiomycetes (24.0%) were dominant, accounting for 78.6% of the total number of fungal sequences obtained (Fig. 1B). Agaricomycetes had its lowest relative abundance at E6 in July. At E1, the relative abundances of Leotiomycetes in May and September were higher than the relative abundance in July, at 40.8% and 47.4%, respectively. The interaction between altitude and season had significant impacts on the relative abundances of Ascomycota and Basidiomycota (*P*<0.05, Table S2).

**Figure 1.**
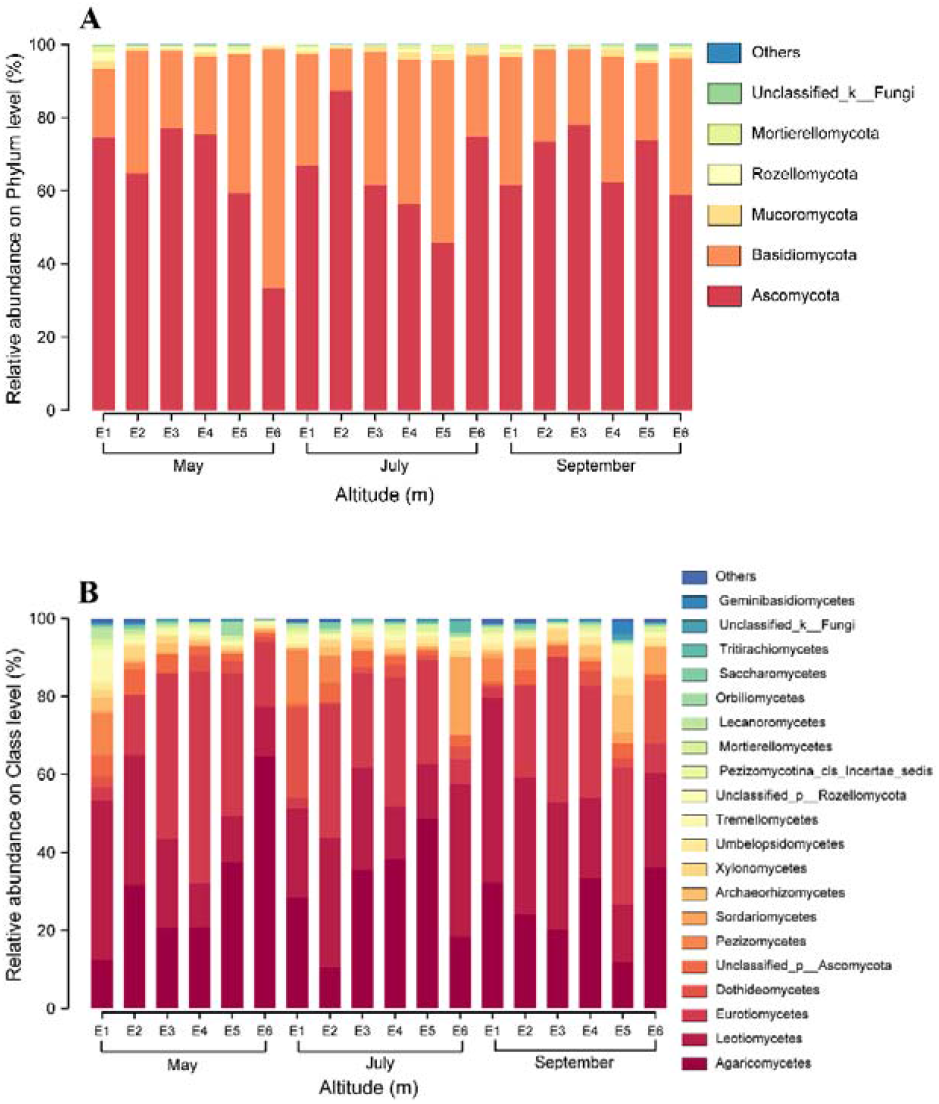
Relative abundances of main soil fungal phyla (A) and classes (B) in different altitudes and seasons soil samples.

### 3.3 Soil fungal community structure and diversity

Altitude had significant effects on the Sobs, Chao1 and Faith’s PD indices of soil fungal diversity (*P*<0.01, Fig. 2). In May and July, the Sobs, Chao1 and Faith’s PD indices were lowest for the highest-altitude (E6) soils among the soils from different altitudes. With increasing elevation, the soil fungal Sobs and Faith’s PD indices showed a decreasing trend (*P*<0.05, Fig. 3). Across May, July, and September, the Sobs index ranged from 373 to 824, the Chao1 index ranged from 553.59 to 1143.56, and Faith’s PD ranged from 74.48 to 153.64. The Simpson index showed significant variation among seasons (*P*<0.05). For the high-altitude soil, this index was highest in May, and in the middle-altitude soils, it was highest in July or September. Altitude and season had significant interaction effects on the Sobs, Chao1 and Faith’s PD indices (*P*<0.05).

**Figure 2.**
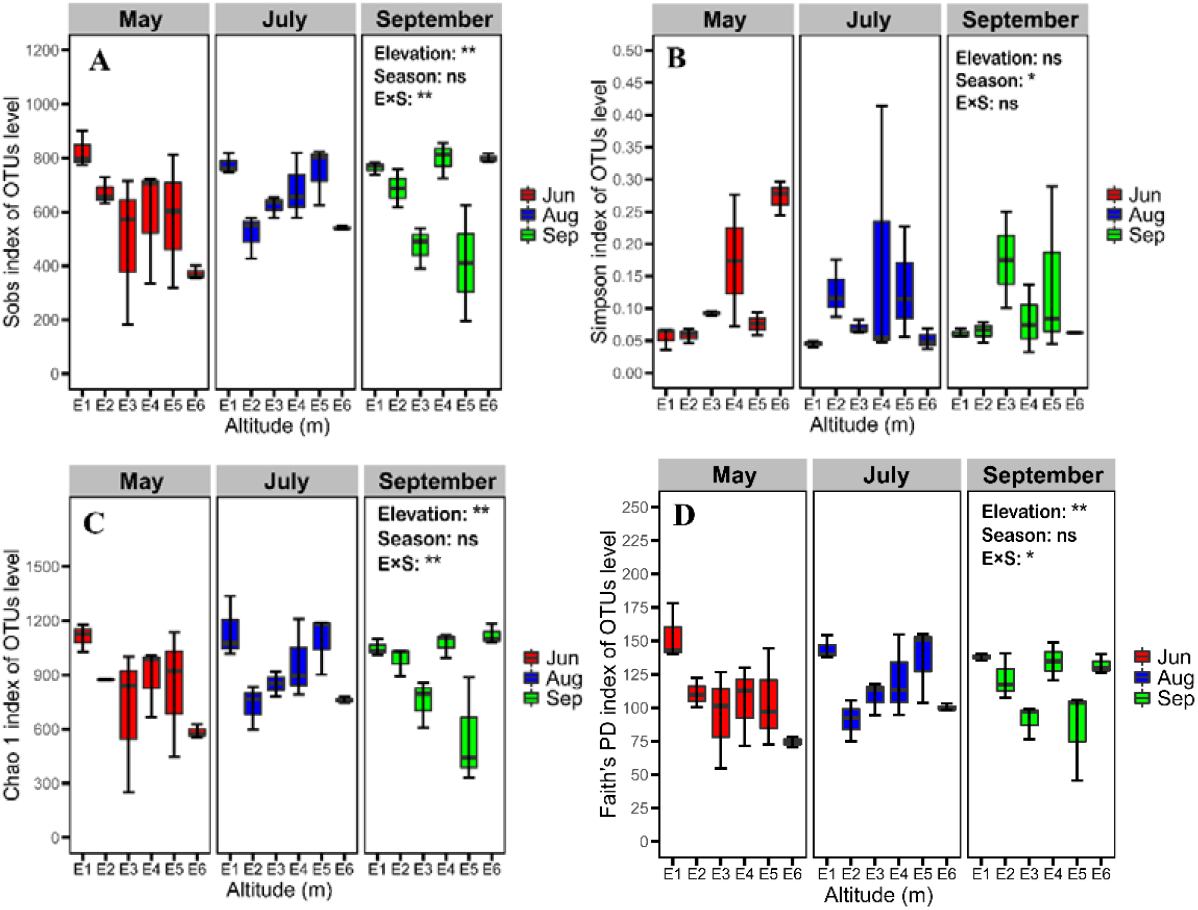
Alpha diversity indices of soil fungal communities in different altitudes soils. OTUs were delineated at 97% sequence similarity. These indices were calculated using random subsamples of 42230 ITS gene sequences per sample. Two-way ANOVA for altitude and season were conducted.

**Figure 3.**
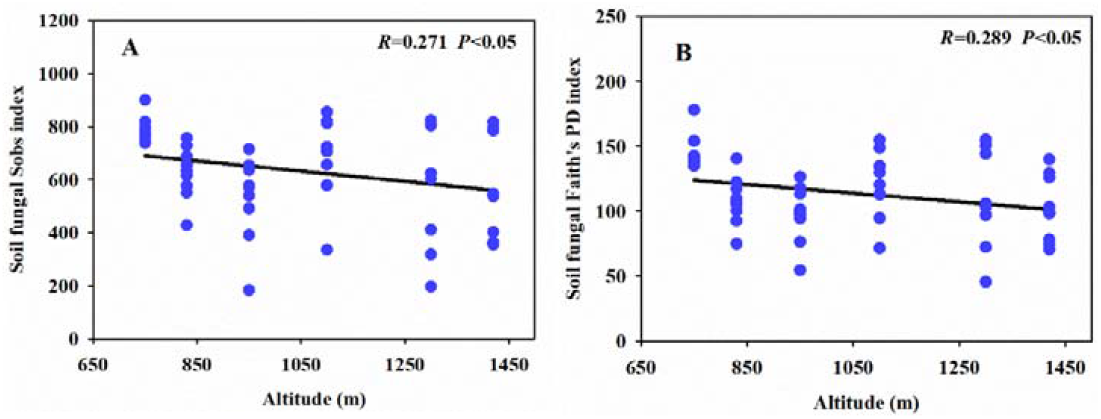
Relationships between soil fungal Sobs, Faith’s PD and altitudes. Diversity indices were calculated using random selections of 42230 sequences per soil sample. The strength of each relationship is based on the linear regression equation.

PCoA based on Bray-Curtis distance was performed on the soil fungal sequencing data corresponding to the different altitudes for each season. In all seasons, the two principal coordinate axes explained more than 30% of the total variation in soil fungi community composition. The samples were divided into three groups by altitude, namely, a low altitude group (E1), a middle altitude group (E2, E3, E4 and E5) and a high altitude group (E6). ANOSIM and PERMANOVA revealed significant differences in the structure of the soil fungal community among altitudes (*P*<0.01, Fig. 4). The PERMANOVA results of all samples demonstrated that altitude had a stronger influence than season on the structure of the soil fungal community (*P*<0.01, Table S3).

**Figure 4.**
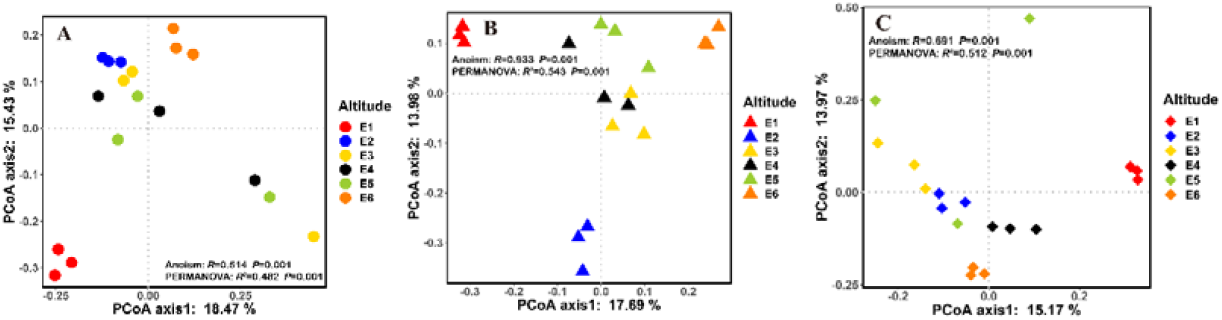
Principal coordinate analysis of soil fungal communities based on Bray-Curtis distances in May (A), July (B) and September (C).

### 3.4 Relationships between soil factors and fungal community structure

The relationships between the soil factors and fungal community structure were evaluated by RDA and the Mantel test. The ordination diagram showed that across all seasons, the first two axes explained more than 50% of the variation in fungal community structure (Fig. 5). However, there were differences among seasons in the main factors affecting community structure. In May, SM was the main influencing factor, and ST and pH played significant roles in July and September, respectively. Other soil factors, such as SOC, TN and DOC, also showed strong effects on fungal community structure (Table 2). Notably, DOC exerted a significant effect on soil fungal community structure in all seasons (Table 2).

**Figure 5.**
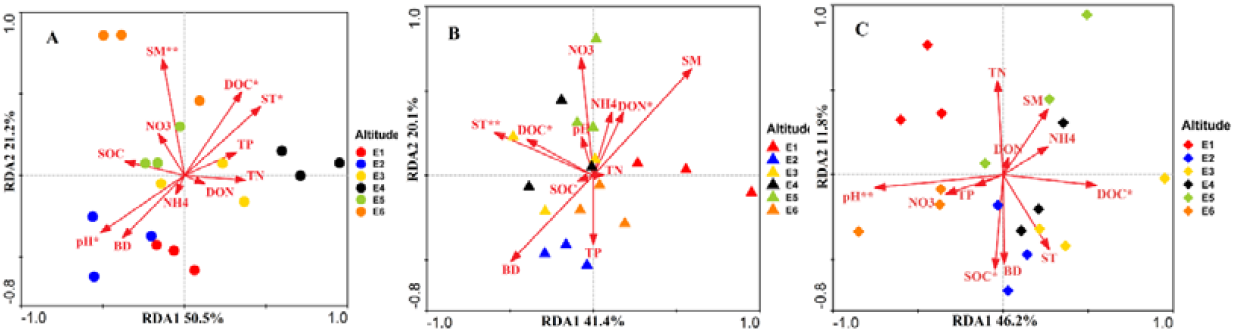
Redundancy analysis based on soil fungal community at genus level (black arrows) and soil factors (red arrows). The top 20 most abundant classified AM fungal genera (97% sequence similarity) in the soil samples. Figure A, B and C represents May, July and September respectively. Direction of arrow indicates the soil factors associated with changes in the community structure, and the length of the arrow indicates the magnitude of the association. The asterisk represents the significant soil factors associated with the fungal community. The percentage of variation explained by RDA 1 and 2 is shown.

**Table 2.** Mantel test results for the correlation between relative abundance of fungal genera and soil variables along the elevational gradient in different seasons.

| Soil variables | May |  | July |  | September |  |
| --- | --- | --- | --- | --- | --- | --- |
| | $R^2$ | $P$ | $R^2$ | $P$ | $R^2$ | $P$ |
| BD | 0.229 | 0.126 | 0.112 | 0.417 | 0.239 | 0.079 |
| SM | 0.404 | **0.005\*\*** | 0.054 | 0.664 | 0.187 | 0.201 |
| ST | 0.314 | **0.033\*** | 0.528 | **0.004\*** | 0.239 | 0.123 |
| TP | 0.099 | 0.456 | 0.067 | 0.610 | 0.027 | 0.808 |
| TN | 0.116 | 0.406 | 0.027 | 0.814 | 0.271 | 0.042 |
| SOC | 0.114 | 0.414 | 0.012 | 0.903 | 0.272 | **0.039\*** |
| NO<sub>3</sub><sup>-</sup>-N | 0.071 | 0.593 | 0.116 | 0.403 | 0.108 | 0.446 |
| NH<sub>4</sub><sup>+</sup>-N | 0.013 | 0.916 | 0.006 | 0.995 | 0.083 | 0.546 |
| pH | 0.312 | **0.039\*** | 0.072 | 0.610 | 0.493 | **0.008\*\*** |
| DOC | 0.300 | **0.037\*** | 0.324 | **0.041\*** | 0.266 | **0.035\*** |
| DON | 0.015 | 0.895 | 0.309 | **0.043\*** | 0.010 | 0.926 |

\* $P < 0.05$ , \*\* $P < 0.01$

### 3.5 Co-occurrence network of soil fungi

Fungal co-occurrence networks were constructed for the different seasons. The OTUs in the network mainly belonged to Ascomycota, Basidiomycota, Mucoromycota, Mortierellomycota, Rozellomycota and unclassified_k_Fungi. Based on a criterion (nodes > 5), the OTUs were divided into eight modules (Figs. 6 and S4). Compared to the networks for May and July, the network for September was larger and more complex. The number of nodes in the September network was 1.5 times that in the July network. In addition, the number of links between nodes was significantly higher and the average connectivity was higher in September than in May and July (Table 3). The network topology in May had a lower average clustering coefficient (avgCC) and higher modularity and average path length (GD) than that in September. A total of 406 links were identified in the September network, including 336 (82.8%) positive links and 70 (17.2%) negative links (Table 3).

**Figure 6.**
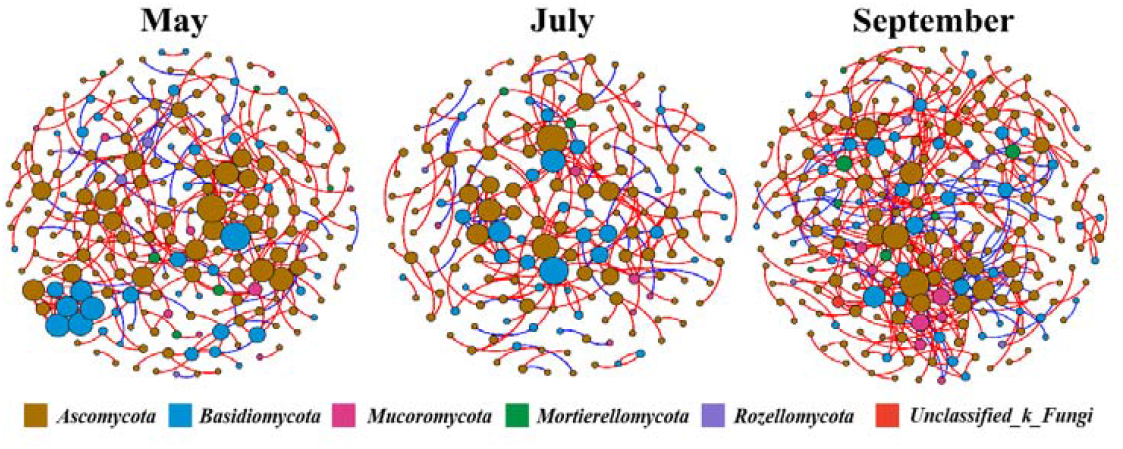
The co-occurrence patterns of soil fungi in different seasons. The size of each node is proportional to the number of degrees. Major phylum (with nodes > 5) were randomly colored. Positive links between nodes were colored red and negative links were colored blue

**Table 3.** The topological properties for soil fungal co-occurrence networks in different seasons

| Network features |  | Season |  |  |
| --- | --- | --- | --- | --- |
|  |  | May | July | September |
| Empirical network | Similarity threshold (St) | 0.790 | 0.790 | 0.790 |
|  | Number of nodes | 221 | 176 | 238 |
|  | Number of links | 278 | 215 | 406 |
| | $R^2$ of power-law | 0.894 | 0.931 | 0.871 |
|  | Number of positive correlations | 237 (85.3%) | 178 (82.8%) | 336 (82.8) |
|  | Number of negative correlations | 41 (14.7%) | 37 (17.2%) | 70 (17.2) |
|  | Average degree (avgK) | 2.516 | 2.443 | 3.412 |
|  | Average clustering coefficient (avgCC) | 0.121 | 0.203 | 0.224 |
|  | Average path distance (GD) | 7.614 | 4.584 | 5.388 |
|  | Modularity | 0.820 | 0.818 | 0.744 |
| Random network | avgCC±SD | 0.010±0.006 | 0.013±0.007 | 0.020±0.007 |
| | GD±SD | 5.104±0.155 | 4.823±0.156 | 4.198±0.061 |
| | Modularity±SD | 0.684±0.011 | 0.682±0.010 | 0.553±0.009 |

Based on the values of *Zi* and *Pi*, we evaluated the possible topological roles of OTUs in the network (Fig. 7). We divided all nodes into four subcategories: peripherals, connectors, module hubs, and network hubs. *Zi-Pi* scatter plots for all nodes in the different seasons were generated based on the module network. No nodes were classified as network hubs, which act as both module hubs and connectors. Most nodes (98.3%) in the fungal network were classified as peripherals, and they were highly connected within their respective modules. Nine nodes belonging to eight genera *(Pezoloma, Cenococcum, Pseudeurotium, Cladophialophora, Archaeorhizomyces*, unclassified_o_GS34, *Pseudogymnoascus*, and *Phialocephala)* were classified as module hubs, and they had strong associations with many nodes in their modules. The nodes of *Meliniomyces* and unclassified_k_Fungi were classified as connectors between modules (Fig. 7). The eleven taxa mentioned above were defined as keystone taxa, indicating that season affected the topological characteristics of individual OTUs and keystone species.

**Figure 7.**
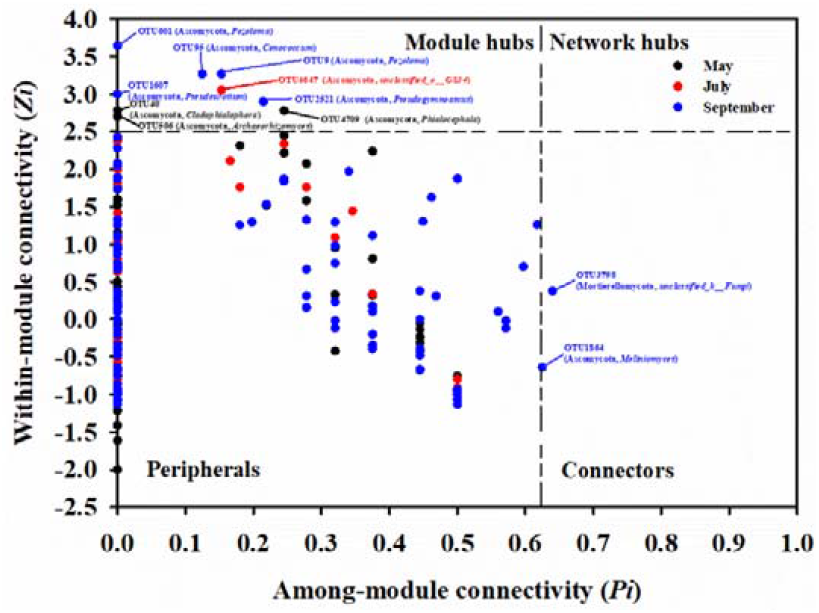
Topological roles of OTUs in the soil fungal co-occurrence networks as indicated by the *Zi-Pi* plot. The nodes with *Zi* > 2.5 are identified as module hubs, and those with *Pi* > 0.62 are connectors. The network hubs are determined by *Zi* > 2.5 and *Pi* > 0.62, and the peripherals are characterized by *Zi* < 2.5 and *Pi* < 0.62

## 4 Discussion

### 4.1 Altitude is more important than season in shaping soil microbial communities in cold-temperate forests

In this study, soil fungal diversity showed a pattern of monotonic decrease along the altitudinal gradient. This result supports our first hypothesis. In addition, ST, SM, pH and DOC played critical roles in influencing fungal diversity. This finding is consistent with previous studies on soil microbial biogeography; numerous studies have demonstrated that most spatial changes in fungal community structure along altitudinal gradients are caused mainly by changes in environmental factors (ST, organic matter content) and vegetative community composition (Bahram et al., 2012; Lugo et al., 2008; Sheng et al., 2019). For instance, soil pH influenced the relative abundances of fungal taxa along an altitudinal gradient on Changbai Mountain (Shen et al., 2014), and environmental factors were significantly associated with the diversity of ectomycorrhizal fungi in the Hyrcanian forest in northern Iran (Bahram et al., 2012). The Mantel test showed that DOC content explained approximately 30% of the variation in fungal community structure in each month, and DOC content increased significantly with increasing elevation. This variation in DOC with elevation may have been caused by the differences in vegetation types among altitudes. Litter inputs from vegetation affect the composition of soil organic matter, which in turn regulates the soil microclimate (Knelman et al., 2012; Shen et al., 2016). Indeed, compared to low-altitude forest soils, alpine tundra soils have more variable organic carbon contents and higher microbial activity (Sjogersten et al., 2003). Shen et al. (2016) reported that DOC was the key factor predicting microbial functional gene diversity along an altitudinal gradient on Changbai Mountain. Nevertheless, in the present study, SM, ST and pH were also important driving factors in May, July and September, respectively. The changes in temperature and precipitation with increasing elevation have direct effects on the fungal community and elicit rapid responses; temperature limits the distribution and physiological activity of animals and plants to some extent (Hodkinson, 2005). Temperature directly affects the fungal community by affecting enzymatic processes and indirectly shapes fungal community structure via its effects on the host plant community and soil nutrients (Hodkinson, 2005). In a study at high latitude, Bahram et al. (2012) found that temperature and precipitation along an altitudinal gradient partially explained the variations in the abundances of ectomycorrhizal fungal groups and fungal community structure. Wang et al. (2015) found that soil pH and temperature drove the variations in soil fungal richness and phylodiversity on Mt. Shegyla on the Tibetan Plateau.

Although several studies have indicated that the large temporal fluctuations of soil microbial community structure observed at the local scale are determined by variations in environmental factors and soil factors among seasons (Diniandreote et al., 2014; Lazzaro et al., 2015; Shigyo et al., 2019), few studies have focused on the seasonality of the soil microbial community at high latitudes. To the best of our knowledge, this is the first report regarding seasonal variations in the altitudinal patterns of soil fungi at high latitudes. Our results do not support our second hypothesis, as the composition (at the phylum and class levels) and structure (as evaluated by PERMANOVA) of the soil fungal community were strongly affected by altitude. In a recent study, Zhang et al. (2020) stated that the influence of spatial heterogeneity on soil microbial communities is more important than the influence of season; they found that compared with bacterial communities, fungal communities were less sensitive to seasonal variation and more sensitive to spatial variation. Fungal communities exhibit slow turnover, preferring low-nutrient, recalcitrant organic matter with a high carbon-nitrogen ratio, and a long substrate cycling time (Blagodatskaya and Anderson, 1998). The size, evolutionary history and niche of individual fungi affect whether they can tolerate large changes in the environment (Schmidt et al., 2014; Sun et al., 2017).

In this study, we found significant seasonal differences in the relative abundances of certain dominant fungi but no significant difference in soil fungal α or β diversity among the different seasons. These findings are in agreement with the findings of Siles and Margesin (2017), who found no obvious seasonal differences in the structure or diversity of soil fungal and bacterial communities at high latitudes; this finding might be related to the absence of significant seasonal variations in key rapidly changing environmental variables. In our study, season had no significant effect on DOC, DON or SM. Zhang et al. (2020) showed that rapidly changing environmental variables (climate factors, SM, and available nutrients) may be the main factors leading to seasonal changes in bacterial and fungal β diversity. In fact, changes in fungal turnover over time cannot be explained only by environmental variations; biotic interactions (such as competition, symbiosis, and predation) and ecological processes also play roles and need be taken into account (Averill et al., 2019). Our results support the finding of Oliverio et al. (2017) that not all taxa in the microbial community are equally sensitive to changes in the environment over time. In the high-latitude boreal forest ecosystem of the present study, although some soil fungal communities showed obvious seasonal variation, altitude had a stronger effect than season on fungal community structure and diversity.

### 4.2 Co-occurrence patterns of soil fungi during seasonal succession

Understanding the interactions among microbial taxa can reveal the spatiotemporal variations in complex microbial community structures (Barberán et al., 2012). Microbial communities with more complex co-occurrence networks are considered to be more resistant to environmental stresses than are those with simpler networks (Banerjee et al., 2019). The network analysis in this study showed that the co-occurrence patterns of the high-latitude fungal communities at different altitudes were nonrandom and that season had a significant impact on the co-occurrence networks of the soil fungal communities. These findings supported our third hypothesis that the co-occurrence networks of soil fungi and their topological characteristics show obvious seasonality. To the best of our knowledge, this is the first study to compare microbial networks among seasons along an altitudinal gradient. Notably, the number of positive links in the network for each season accounted for more than 80% of the total links, which indicates that mutualism or commensalism may play a key role in shaping the fungal community structure at high latitudes. However, negative links were also present in the networks, which indicates that competitive interactions between soil fungi may also occur (Chen et al., 2019). Based on the RMT, at the same *St*, number of network nodes and links, the average degree (avgK) and avgCC were higher in September than in May and July. The higher avgCC in September indicated that the number of within-cluster links was higher than the number of between-cluster links, indicating that soil fungi in similar niches were more closely linked than were those in dissimilar niches. Compared with the September network, the May and July networks were more obviously modular. In a highly modular species network, fungal community structure is largely stable and ordered, and fungi can exchange energy efficiently (Lu et al., 2013).

Keystone species play vital roles in driving the structure and function of microbial communities (Banerjee et al., 2018; Deng et al., 2012). Ma et al. (2016) indicated that, based on network growth processes, key nodes are considered the initial components of the network assembly. In our study, Ascomycota and Basidiomycota played dominant roles in the network, and the nine OTU nodes that were module hubs belonged to Ascomycota *(Cladophialophora, Archaeorhizomyces, Phialocephala*, unclassified_o_GS34, *Pseudeurotium, Pseudogymnoascus, Pezoloma*, and *Cenococcum).* Members of *Cladophialophora* are saprophytic nutrient fungi, and *Archaeorhizomyces* and *Phialocephala* can improve the availability of soil phosphorus, which in turn promotes plant growth and development (Jumpponen et al., 1998; Zhang et al., 2018). Due to our poor understanding of some fungal functions, it is impossible to fully infer the roles of these keystone taxa in the environment. Nevertheless, the positive correlations among these ascomycete taxa are consistent with previous findings (Barberán et al., 2012; Faust and Raes, 2012), which suggests that these ascomycetes may act synergistically in the environment or share similar niches. Overall, the present study showed that season had little effect on fungal diversity and community structure along the altitudinal gradient, that the soil fungal co-occurrence network exhibited obvious seasonal succession, and that environmental changes caused by seasonal changes regulated the symbiotic networks and keystone species.

## 5 Conclusions

In summary, soil fungal diversity showed a pattern of monotonic decrease along an altitudinal gradient in a high-latitude region. Soil moisture, temperature, pH and DOC were the main factors affecting the distribution of soil fungi in the different seasons. We found that although altitude had stronger effects than season on fungal community structure and fungal diversity, the soil fungal co-occurrence network exhibited obvious seasonal succession, indicating that the keystone species of soil fungi exhibited niche differentiation among seasons. These results emphasize the importance of time-scale sampling of soil fungal communities to better understand the succession and assembly processes of soil microbial communities in natural ecosystems.

## Authors’ contributions

L.J. and L.Y. conceived and designed the original conceptual idea. L.J. and Y.Z. performed experimental work, sampling and DNA sequencing. L.J. carried out the bioinformatics and statistical analysis. L.J. created the figures and wrote the manuscript. L.J., Y.Y. and L.Y. provided fundings. All authors helped to edit and complete the manuscript.

## Acknowledgements

This work was financially supported by the National Key Research and Development Program of China (2017YFD0601204), the Fundamental Research Funds for the Central Universities (2572019AA07; 2572019CP16), and the Heilongjiang Touyan Innovation Team Program (Technology Development Team for Highly efficient Silviculture of Forest Resources). We thank Lixin Ma, Qingchao Zhu and the A’longshan Forestry Bureau for access permission and logistic support. We also thank Jiangbo Yu and Yujiao Wang for assistance in laboratory analyses, and Guigang Lin for their constructive comments.

