## Supplemental files for "Seasonal variations in soil fungal communities and co-occurrence networks along an altitudinal gradient in the cold temperate zone of China: A case study on Oakley Mountain"

Supplementary material captions

**Supplementary Table 1** Site characteristics of different altitudes

**Supplementary Table 2** Two-way analysis of variance of relative abundance of main fungal phyla and class

**Supplementary Table 3** Permutation multivariate analysis of variance (PERMANOVA) of soil fungal community between altitude and season

**Supplementary Figure 1** Location of the Oakley Mountains in Greater Khingan Mountains

**Supplementary Figure 2** Average temperature and precipitation in the sampling month

**Supplementary Figure 3** The Rarefaction curves of the number of operational taxonomic units (OTUs) for soil fungal communities in May (A), July (B) and September (C). Random subsamples of 42230 ITS gene per sample were used to generate the rarefaction curves. OTUs were delineated at 97% sequence similarity.

**Supplementary Figure 4** The co-occurrence patterns of soil fungi in different seasons.

The size of each node is proportional to the number of degrees. Major modules (with nodes > 5) were randomly colored. Positive links between nodes were colored red and negative links were colored blue

**Supplementary Table 1** Site characteristics of different altitudes

| Site | Coordinates | Altitude  (ma.s.l.) | Soil type | Vegetation type | Dominant taxa |
| --- | --- | --- | --- | --- | --- |
| E1 | N 51°45′52″  E 122°06′19″ | 750 | Grass brown coniferous forest soil | Cold temperate coniferous forest | *Betula platyphylla*、*Chosenia arbutifolia*、*Larix gmelinii*、*Rhododendron lapponicum*、*Ledum palustre*、*Vaccinium vitis-idaea*、*Alnus sibirica*、*Swida alba*、*Malus baccata*、*Padus racemosa*、*Deyeuxia angustifolia*、*Equisetum pratense* |
| E2 | N 51°47′41″  E 122°05′03″ | 830 | Dark brown coniferous soil | Cold temperate coniferous forest | *Larix gmelinii*、*Vaccinium vitis-idaea*、*Ledum palustre*、*Rosa davurica*、*Lonicera caerulea*、*Rubus arcticus*、*Pyrola incarnata*、*Deyeuxia angustifolia* |
| E3 | N 51°49′42″  E 122°03′34″ | 950 | Dark brown coniferous soil | Cold temperate coniferous forest | *Larix gmelinii*、*Betula platyphylla*、*Vaccinium vitis-idaea*、*Ledum palustre*、*Rhododendron dauricum*、*Spiraea dahurica*、*Rubus sachalinensis*、*Sambucus williamsii*、*Rosa davurica*、*Artemisia lagocephala*、*Cimicifuga foetida*、*Vicia ramuliflora*、*Pyrola incarnata* |
| E4 | N 51°49′59″  E 122°02′46″ | 1100 | Podzolic brown coniferous soil | Cold temperate coniferous forest | *Larix gmelinii*、*Pinus sylvestris*、*Pinus pumila*、*Betula platyphylla*、*Vaccinium vitis-idaea*、*Ledum palustre*、*Rhododendron dauricum*、*Spiraea dahurica*、*Rubus sachalinensis*、*Sambucus williamsii*、*Artemisia lagocephala*、*Clematis sibirica*、*Cimicifuga foetida*、*Vicia ramuliflora* |
| E5 | N 51°50′14″  E 122°02′19″ | 1300 | Podzolic brown coniferous soil | Cold temperate coniferous forest | *Larix gmelinii*、*Pinus pumila*、*Betula ermanii*、*Rhododendron dauricum*、*Vaccinium vitis-idaea*、*Ledum palustre*、*Artemisia lagocephala*、*Aquilegia viridiflora*、*Saxifraga bronchialis*、*Polygonum alpinum* |
| E6 | N 51°51′24″  E 122°02′7″ | 1420 | Podzolic brown coniferous soil | Subalpine dwarf forest | *Pinus pumila* |

m a.s.l. = meters above sea level

**Supplementary Table 2** Two-way analysis of variance of relative abundance of main fungal phyla and class

| Taxonomy | | Altitude | | Season | | Altitude×Season | |
| --- | --- | --- | --- | --- | --- | --- | --- |
|  |  | *F* | *P* | *F* | *P* | *F* | *P* |
| Phyla | Ascomycota | 2.124 | 0.085 | 0.316 | 0.731 | 2.474 | **0.023** |
|  | Basidiomycota | 2.049 | 0.095 | 0.406 | 0.669 | 3.027 | **0.007** |
|  | Mucoromycota | 3.293 | **0.015** | 2.603 | 0.088 | 1.979 | 0.066 |
|  | Rozellomycota | 2.069 | 0.092 | 0.774 | 0.469 | 0.913 | 0.532 |
|  | Mortierellomycota | 3.127 | **0.019** | 0.118 | 0.889 | 1.436 | 0.204 |
|  | Unclassified_k__Fungi | 0.954 | 0.458 | 1.401 | 0.259 | 0.648 | 0.763 |
|  | Others | 1.596 | 0.186 | 0.472 | 0.628 | 0.683 | 0.733 |
| Classes | Agaricomycetes | 2.069 | 0.092 | 0.664 | 0.521 | 4.018 | **0.001** |
|  | Leotiomycetes | 13.485 | **<0.001** | 3.788 | **0.032** | 3.107 | **0.006** |
|  | Eurotiomycetes | 6.108 | **<0.001** | 0.798 | 0.458 | 0.677 | 0.738 |
|  | Dothideomycetes | 28.942 | **<0.001** | 14.945 | **<0.001** | 32.638 | **<0.001** |
|  | Unclassified_p__Ascomycota | 3.739 | **0.008** | 1.663 | 0.204 | 1.735 | 0.110 |
|  | Pezizomycetes | 41.660 | **<0.001** | 3.727 | 0.034 | 2.894 | **0.009** |
|  | Sordariomycetes | 24.529 | **<0.001** | 9.783 | **<0.001** | 10.457 | **<0.001** |
|  | Archaeorhizomycetes | 0.739 | 0.599 | 0.911 | 0.411 | 1.453 | 0.198 |
|  | Xylonomycetes | 3.032 | **0.022** | 1.538 | 0.229 | 2.534 | **0.020** |
|  | Umbelopsidomycetes | 3.680 | **0.009** | 2.956 | 0.065 | 2.019 | 0.060 |
|  | Tremellomycetes | 2.139 | 0.083 | 0.751 | 0.479 | 1.603 | 0.145 |
|  | Unclassified_p__Rozellomycota | 1.976 | 0.106 | 0.855 | 0.434 | 0.943 | 0.507 |
|  | Pezizomycotina_cls_Incertae_sedis | 11.034 | **<0.001** | 4.270 | **0.022** | 2.677 | **0.015** |
|  | Mortierellomycetes | 3.127 | **0.019** | 0.118 | 0.889 | 1.435 | 0.205 |
|  | Lecanoromycetes | 36.662 | **<0.001** | 35.736 | **<0.001** | 14.760 | **<0.001** |
|  | Orbiliomycetes | 1.223 | 0.318 | 0.698 | 0.504 | 0.862 | 0.576 |
|  | Saccharomycetes | 7.395 | **<0.001** | 0.993 | 0.381 | 0.453 | 0.909 |
|  | Tritirachiomycetes | 21.496 | **<0.001** | 34.837 | **<0.001** | 17.853 | **<0.001** |
|  | Unclassified_k__Fungi | 0.954 | 0.458 | 1.401 | 0.259 | 0.648 | 0.763 |
|  | Geminibasidiomycetes | 0.957 | 0.457 | 0.860 | 0.432 | 1.055 | 0.420 |
|  | Others | 8.627 | **<0.001** | 3.868 | **0.030** | 0.945 | 0.505 |

**Supplementary Table 3** Permutation multivariate analysis of variance (PERMANOVA) of soil fungal community between altitude and season

|  |  | Df | Sums of Sqs | Mean Sqs | F.Model | *R^2^* | Pr(>F) |
| --- | --- | --- | --- | --- | --- | --- | --- |
| Bray-Curtis | Altitude | 5 | 6.967 | 1.393 | 7.667 | **0.444** | **0.002** |
|  | Season | 2 | 0.784 | 0.392 | 1.342 | 0.050 | 0.111 |

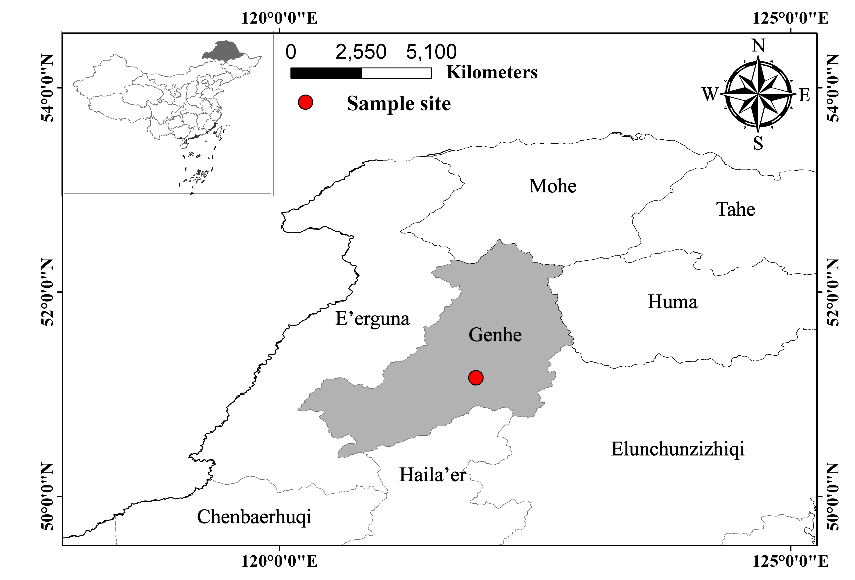

**Supplementary Figure 1** Location of the Oakley Mountains in Greater Khingan Mountains

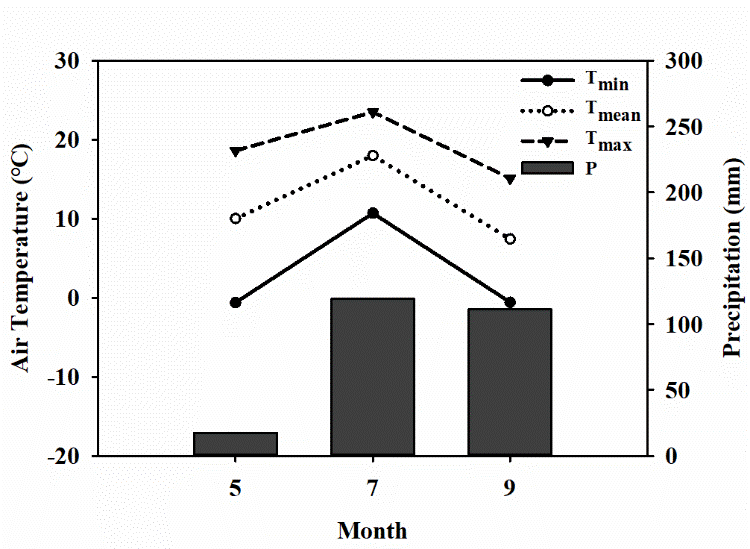

**Supplementary Figure 2** Average temperature and precipitation in the sampling month

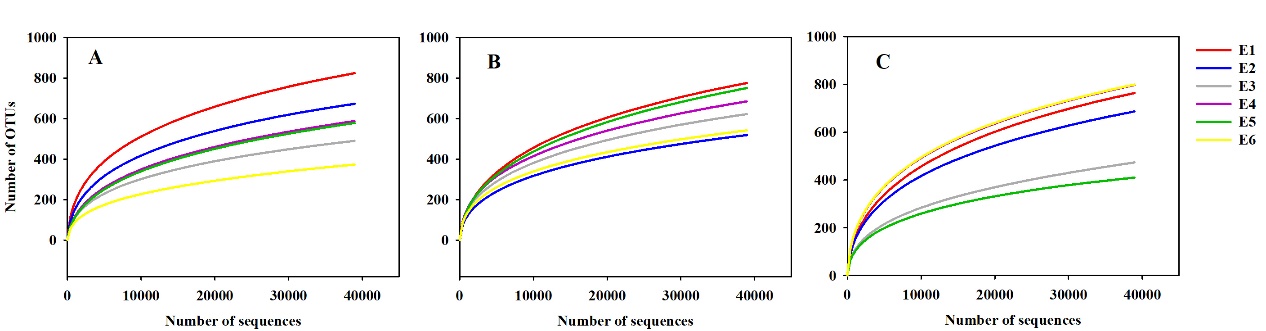

**Supplementary Figure 3** The Rarefaction curves of the number of operational taxonomic units (OTUs) for soil fungal communities in May (A), July (B) and September (C). Random subsamples of 42230 ITS gene per sample were used to generate the rarefaction curves. OTUs were delineated at 97% sequence similarity.

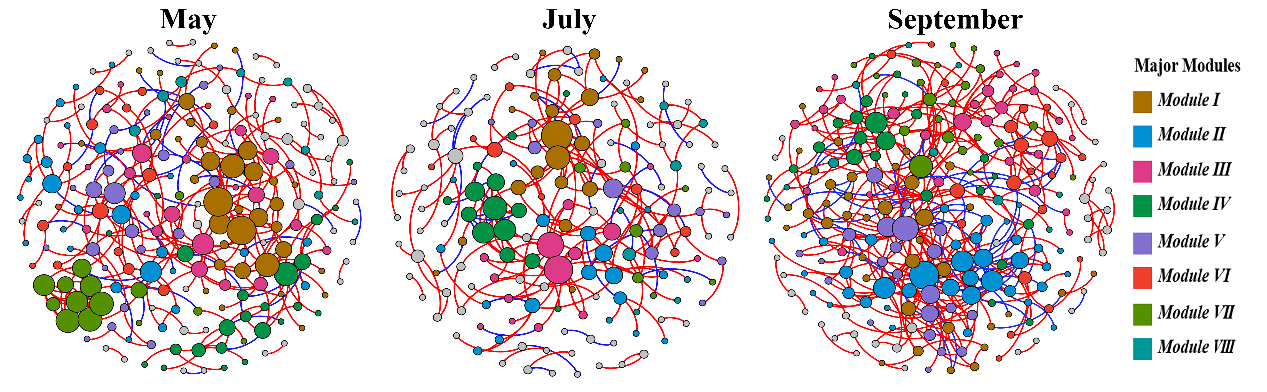

**Supplementary Figure 4** The co-occurrence patterns of soil fungi in different seasons.

The size of each node is proportional to the number of degrees. Major modules (with nodes > 5) were randomly colored. Positive links between nodes were colored red and negative links were colored blue
